# The Tubulin Nano-Code: a protofilament-specific pattern of tubulin post-translational modifications regulates ciliary beating mechanics

**DOI:** 10.1101/2023.06.28.546853

**Authors:** G. Alvarez Viar, N. Klena, F. Martino, A. Nievergelt, G. Pigino

**Affiliations:** Human Technopole, Milan, Italy; Max Planck Institute of Molecular Cell Biology and Genetics, Dresden, Germany

## Abstract

Control of ciliary beating is crucial for motility and signaling in eukaryotic cells and requires spatially restricted interactions between axonemal proteins and specific protofilaments within the ciliary microtubules. How these interactions are regulated remains poorly understood, but increasing evidence indicates that tubulin post-translational modifications (tPTMs) are required for proper ciliary motility. The Tubulin Code refers to the concept that tPTMs can modulate the function of individual microtubules in cells. Here we use a combination of immuno-cryo-electron tomography, expansion microscopy and mutant analysis to show that, in motile cilia, tubulin glycylation and polyglutamylation form mutually exclusive protofilament-specific nano-patterns at sub-microtubular scale. We show that these two nano-patterns are consistent with the distributions of axonemal dyneins and nexin-dynein regulatory complexes, respectively, and are required for their regulation during ciliary beating. Our discovery of a tubulin nano-code in cilia highlights the need of higher-resolution studies also in other cellular compartments to understand the molecular role of tPTMs.

**One-Sentence Summary:** Tubulin post-translational modifications form nanopatterns at sub-microtubular scale that enable individual protofilaments to perform specific functions

**Graphical Abstract:** 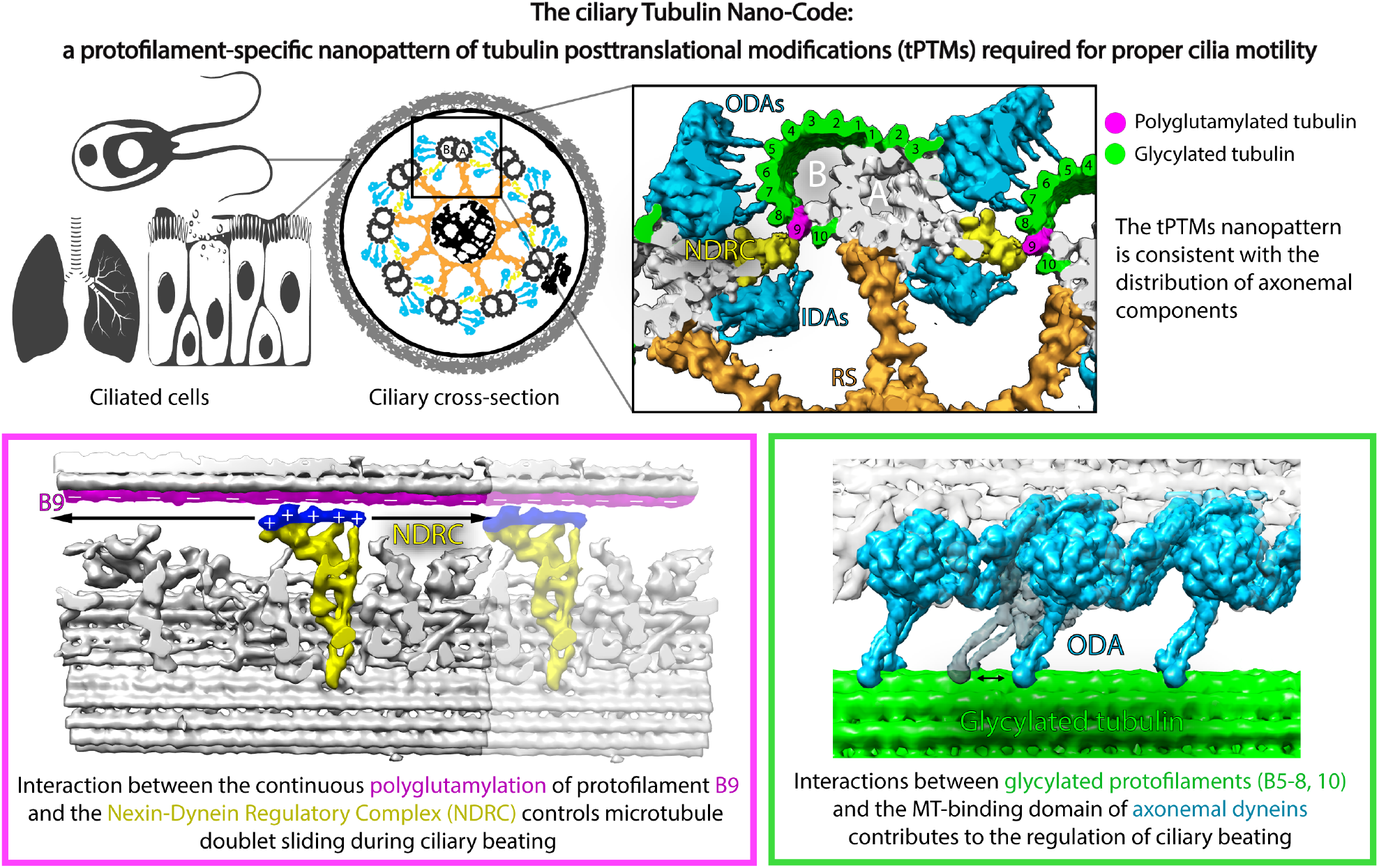

## Main Text

Cilia are thin, elongated, microtubule-based organelles with key conserved roles in organismal development and homeostasis (1-4). Their correct function relies on the precise organization of different macromolecular components along a microtubular scaffold called the axoneme (5-10). Axonemal protein complexes, such as dynein arms and the nexin-dynein regulatory complex (NDRC) stably bind on the A-tubule of the microtubule doublet, and transiently interact with the B-tubule of the adjacent microtubule doublet to perform their functions. Both stable and transient interactions form repetitive patterns along the axoneme. For instance, outer dynein arms (ODAs) bind to the A-tubule on protofilaments A3 and 4 every 24 nm and interact with protofilaments B5-8 of the adjacent B-tubule with their microtubule binding domains, while NDRC binds to A7-9 every 96 nm and interacts with protofilament B9 of the neighboring B-tubule (11-13) (Figure 1 A). Historically, microtubules were considered mere structural components of the axoneme with no regulatory function in ciliary beating. However, cilia are highly enriched in tPTMs, which can functionally differentiate subpopulations of microtubules or individual microtubules in the cell (reviewed in 14). Given the ubiquitous interaction of different axonemal proteins with microtubules, tPTMs might be expected to contribute to the assembly and/or function of axonemal and ciliary components. Although tPTMs have been shown to be required for proper ciliary function (15-21), however, the molecular mechanisms remain elusive.

**Figure 1:**
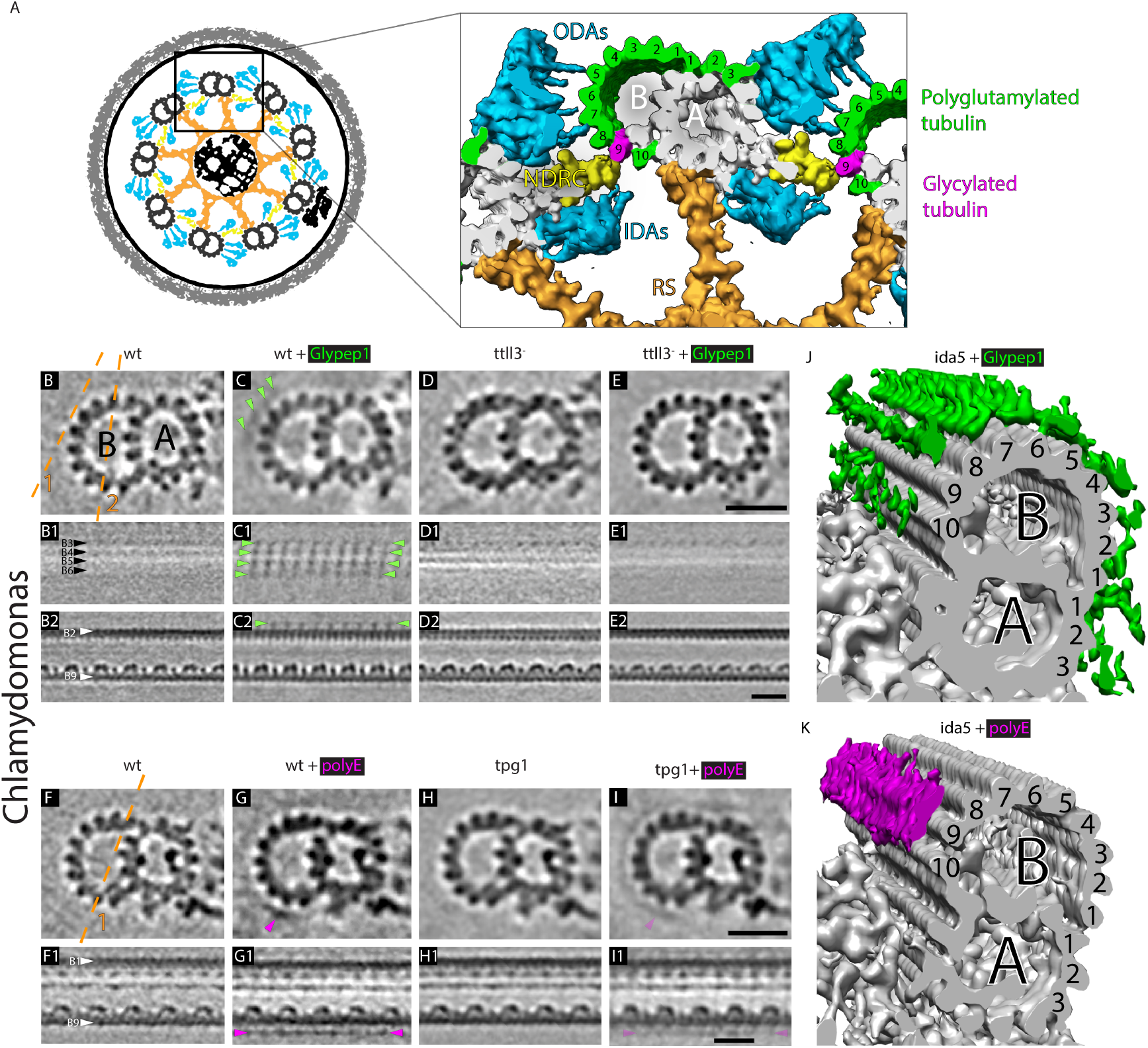
Localization of tubulin glycylation and polyglutamylation in the Chlamydomonas axoneme. **A:** Cartoon representing the ciliary cross section and a close up view of the 96-nm repeat showing the spatially restricted interactions between axonemal components and different protofilaments of the adjacent doublet. **B to I**: Slices through 3D electron density models of the Chlamydomonas axonemal 96-nm repeat generated by sub-tomogram averaging. Dashed lines 1 and 2 in **B** depict the planes at which the different longitudinal sections were made. **B**: wild type axoneme. **C**: wild type axoneme decorated with Glypep1 antibodies. Note the presence of extra densities, indicated by green arrowheads, over the microtubule doublet surface recapitulating the tubulin dimer periodicity, indicating the presence of the antibody labeling and the position of glycylated tubulin. **D**: *ttll3^-^* axoneme. **E**: *ttll3^-^* axoneme incubated with Glypep1 antibodies. Note the absence of any labeling. **F**: wild type axoneme. **G**: wild type axoneme decorated with polyE antibodies. Note the presence of extra densities, indicated by magenta arrowheads, present over protofilament B9. **H**: *tpg1* axoneme. **I**: *tpg1* axonemes incubated with polyE antibodies. Note the reduction in the labeling signal compared to G. **J**: Electron density model of the *ida5* 96-nm repeat decorated with Glypep1 antibodies (green). **K**: Electron density model of the *ida5* 96-nm repeat decorated with polyE antibodies (magenta). A and B labels indicate the different tubules in the microtubule doublet. Black and white arrowhead labels indicate different protofilaments. Scale bars 20nm.

Work from our lab has shown that tubulin glycylation is required for proper ciliary beating in mouse sperm, as it contributes to the regulation of axonemal dynein activity (22). However, it remains unclear whether this is a consequence of a direct physical interaction between the microtubule binding domains of axonemal dyneins and glycylated tubulin sites. We hypothesize that other tPTMs might be involved in the specific regulation of other axonemal components. To achieve such a spatially constrained regulatory function, tPTMs would have to be distributed over the microtubule surface forming a pattern at the sub-microtubular scale that is consistent with the distribution of functional interactions between axonemal components and individual protofilaments.

Previous studies that have investigated the distribution of tPTMs in the axoneme of Chlamydomonas showed differences in modifications between the central pair and microtubule doublets, and even between the A-tubule and the B-tubule of each doublet (23-25). However, these studies were limited in resolution, and did not explore the putative regulatory role of tPTMs on the function of different axonemal protein complexes. In this study, we address this issue by resolving the distribution of two tPTMs at the nanoscale and test our hypothesis of the presence of a tubulin code nanopattern that would reflect the axonemal architecture. Our results demonstrate that a conserved nanopattern of polyglutamylation and glycylation in the axoneme is required for the proper function of the NDRC and axonemal dyneins, respectively, thus shedding light on the role of tPTMs in ciliary mechanics.

## Results

### Tubulin glycylation and polyglutamylation form a complementary protofilament-specific pattern over the axonemal microtubule doublet in Chlamydomonas

To test our hypothesis that axonemal tPTMs distribution reflects the organization of the axoneme, we developed a protocol to visualize their localization at the nanoscale. We incubated axonemes with antibodies against different tPTMs, plunge-froze these samples and performed cryo-electron tomography and sub-tomogram averaging (Supplementary Figure 1 A, see Material and Methods for details). This approach resolved the binding site of antibodies against different tPTMs in electron density models of the axonemal 96-nm repeat, revealing the location of specific tPTMs in flagella/cilia of Chlamydomonas cells. Wild type control axonemes (without antibodies) did not show any decoration over the exposed surface of the microtubule doublet (Figure 1 B). Upon addition of Glypep1 antibodies, against glycylated tubulin, we observed obvious extra densities over most of the exposed microtubule surface, comprising protofilaments A1-3, B1-8 and B10 (Figure 1 C). The antibody decoration clearly followed the tubulin dimer repeat (Figure 1 C1), reflecting the expected antigenicity of the antibody against glycylated beta-tubulin.

As a negative control to validate the specificity of the antibody, we analyzed axonemes from the glycylation knockout strain ttll3- (CLiP library, strain 124984 (26), Supplementary Figure 1 B). The structure of the 96-nm repeat, with the typical microtubule-associated axonemal complexes, appeared identical between the ttll3-strain and wild type (Figure 1 D, Supplementary Figure 2 A and B). As expected, upon addition of Glypep1 antibodies to ttll3-axonemes, no decoration was found (Figure 1 E). This data shows that glycylation occupies protofilaments A1-3 and most of the B-tubule surface, with the exception of protofilament B9 (Figure 1 C2 and J). Interestingly, the microtubule binding domains for inner and outer dynein arms (IDAs and ODAs, respectively) interact with protofilaments of the B-tubule (B5-10) (11-13). Thus, the microtubule binding domains of axonemal dyneins interact directly with tubulin glycylation. Consistent with this, the absence of glycylation in mouse sperm axonemes causes defective ODA and IDA activity and an impaired motility phenotype (22).

Tubulin glycylation and polyglutamylation are thought to compete for the same initiation sites, and tubulin polyglutamylation is predicted to be located preferentially on the B-tubule (24, 25, 27). To investigate the localization of polyglutamylation, we incubated isolated axonemes with polyE antibodies raised against polyglutamate chains and observed a clear enrichment along protofilament B9 (Figure 1D-G). As a negative control for the localization of the polyE antibody, we used the strain tpg1, whose levels of axonemal tubulin polyglutamylation are significantly decreased as a result of knocking out the polyglutamylase TTLL9 (19). Analogous to the glycylation KO mutant (ttll3-), the axonemal 96-nm repeat structure of tpg1 appeared normal (Figure 1 H, Supplementary Figure 2 A and C). Upon addition of polyE antibodies, tpg1 showed a much reduced signal on B9 (Figure 1 I), consistent with the phenotype of this strain. Protofilament B9 is near to the IDA microtubule binding domains and stalks, which are often visible in the averages of wild type axonemes. Because of the positioning of the polyglutamylation associated signal over protofilament B9, we wanted to exclude the potential contribution of these binding domains and stalks. Therefore, we used the strain ida5, which lacks most IDAs, and immunolabeled its axonemes with polyE and Glypep1 antibodies separately. Imaging reinforced the results obtained with wild type axonemes (Supplementary Figure 3), where tubulin polyglutamylation is enriched over protofilament B9 and glycylation occupies the rest of the exposed microtubular surface.

Taken together, these results demonstrate that it is possible to precisely locate tPTMs at nanometric level on native microtubular assemblies and revealed that tubulin glycylation and polyglutamylation form a complementary protofilament-specific pattern in the Chlamydomonas axoneme (Figure 1 A, J and K).

### The tubulin glycylation and polyglutamylation nanopattern is conserved in motile cilia from algae to mammals

To test the conservation of this pattern in motile cilia of other eukaryotes, we isolated mouse trachea respiratory cilia and performed the same immuno-cryo-ET procedure. Analysis of control axonemes (without antibodies) showed the typical structure of the 96-nm repeat of mouse respiratory cilia (28) (Figure 2 A, Supplementary Figure 4). Upon addition of Glypep1 antibodies, most of the exposed microtubular surface was decorated by extra densities (protofilaments A1-3, B1-8 and B10) (Figure 2 B,C) with protofilament and repeat patterns that recapitulated what we observed in Chlamydomonas (Figure 1 B1, E). Upon addition of polyE antibodies, a clear decoration was observed specifically over protofilament B9 (Figure 2 E,F). Thus, the complementarity of the glycylation and polyglutamylation nanopatterns is conserved between distantly related species like algae and mice, suggesting it is a characteristic of all motile cilia. Such a conserved pattern calls for a fundamental role of these two tPTMs in the correct function of motile cilia across species.

**Figure 2:**
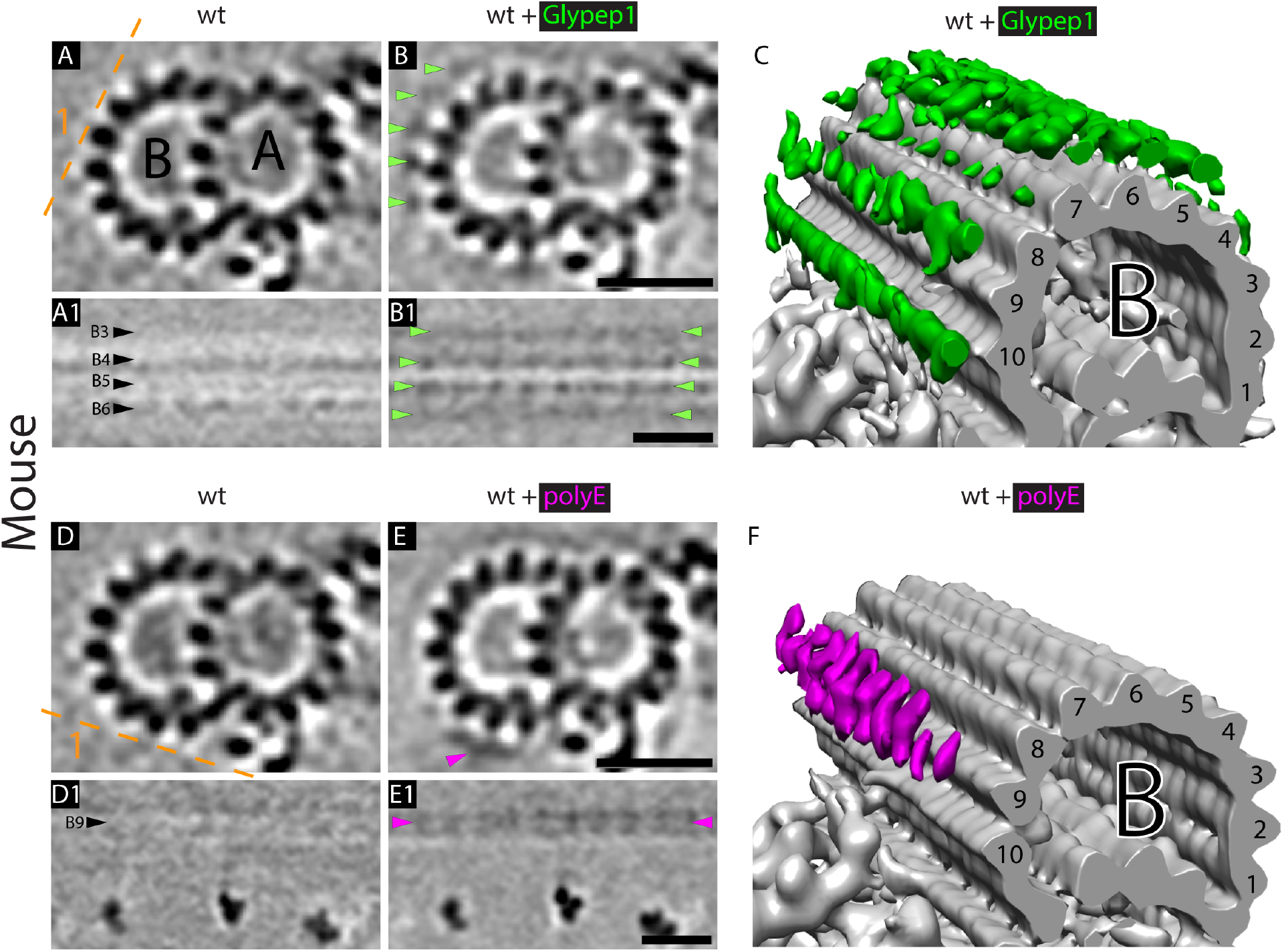
Localization of tubulin glycylation and polyglutamylation in the mouse respiratory axoneme. Slices through 3D electron density models of the mouse respiratory cilia axonemal 96-nm repeat generated by sub-tomogram averaging. Dashed lines depict the planes at which the different longitudinal sections were made. A and B labels indicate the different tubules in the microtubule doublet. Black arrowhead labels indicate different protofilaments. **A**: Mouse respiratory cilia axonemes isolated from trachea tissue. **B**: Mouse respiratory cilia axonemes decorated with Glypep1 antibodies. Note the presence of extra densities, indicated by green arrowheads, over the microtubule doublet surface recapitulating the tubulin dimer periodicity. **C**: Electron density model of the mouse respiratory cilia axoneme decorated with Glypep1 antibodies (green). **D**: Mouse respiratory cilia axonemes isolated from trachea tissue. **E**: Mouse respiratory cilia axonemes decorated with polyE antibodies. Note the presence of extra densities, indicated by magenta arrowheads, over protofilament B9. **F**: Electron density model of the mouse respiratory cilia axoneme decorated with polyE antibodies (magenta). Scale bar: 1um.

### How is the tubulin glycylation and polyglutamylation nanopattern generated?

The enzymes that generate tPTMs are thought to be spatially and temporally regulated to modulate the interaction of proteins with microtubules in different parts of the cell and in different moments of the cell cycle (14). Our results show that modifying enzymes are able to generate a precise nanopattern of glycylation and polyglutamylation at the level of a single protofilament. We then set out to investigate how this tPTMs pattern is formed during ciliary assembly using ultrastructural expansion microscopy of regrowing cilia of Chlamydomonas.

We simultaneously labeled acetylated tubulin, tubulin and either polyglutamylated or glycylated tubulin at different stages of ciliary regeneration (Figure 3). Ciliary tips, where tubulin dimers are added to the plus-ends of A-microtubules, showed an intense signal for tubulin antibodies, while none of the modifications we studied were present in this area (Figure 3 A, C, E1 and E2, Supplementary Figure 4 and 5). These results were consistent in both growing and steady state ciliary tips (Figure 3 A and C, Supplementary Figure 5 and 6). During ciliary growth, the tubulin acetylation signal was located just proximally with respect to the unmodified tip area. Glycylation and polyglutamylation are instead located proximally with respect to acetylation leading edge indicating that acetylation is established before glycylation and polyglutamylation (Figure 3 A and C). The intensity of the fluorescence signal associated with tubulin acetylation along the cilium was constant during ciliary assembly and in full-length cilia. Similarly, axonemal polyglutamylation levels remain constant regardless of the ciliary growth stage (Figure 3 B, Supplementary Figure 5). On the other hand, tubulin glycylation slowly builds up along the axoneme during ciliary growth, showing a clear difference in relative intensity between short (0-5 micron) and long (>5 micron) cilia (Figure 3 D and Supplementary Figure 6). Based on these results, we propose that polyglutamylation enzymes carry out their activity mostly at the tip of newly assembled B-tubules and are able to quickly modify all polyglutamylation targets, while glycylation enzymes take significantly longer time to fully decorate the remaining sites on the B-tubule and A-tubule lattices that we showed by immuno-cryo-ET.

**Figure 3:**
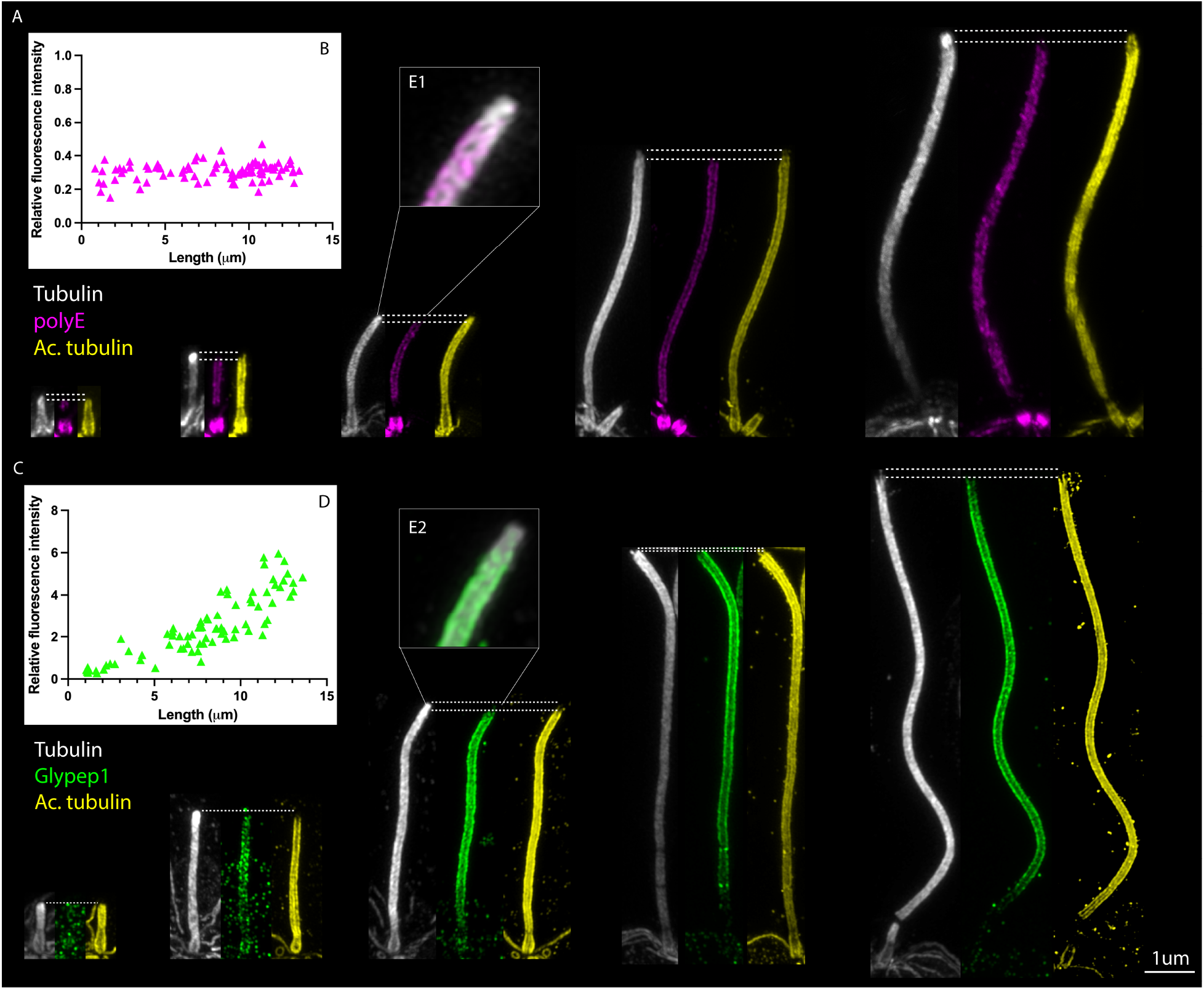
Axonemal incorporation of tubulin polyglutamylation and glycylation during ciliary regrowth in Chlamydomonas. Ultrastructural expansion microscopy of regrowing flagella of Chlamydomonas. **A**: Triple immunostaining of regrowing chlamydomonas axonemes against tubulin (gray), polyglutamylated tubulin (magenta) and acetylated tubulin (yellow). Note that even for very short cilia polyglutamylation is readily incorporated. **B**: Quantification of polyE labeling density over regrowing cilia normalized to polyE signal in the basal body (representative images for this quantification can be found in Supplementary Figure 4). **C**: Triple immunostaining of regrowing chlamydomonas axonemes against tubulin (gray), glycylated tubulin (green) and acetylated tubulin (yellow). Note that for short cilia glycylation signal is sparse while its density increases for long cilia. **D**: Quantification of Glypep1 labeling density over regrowing cilia normalized to centrin signal in the basal body (representative images for this quantification can be found in Supplementary Figure 5). **E1 and 2**: Blow-up from a single ciliary tip showing the overhang of the tubulin signal with respect to the respective tPTMs

### Lack of glycylation affects phototaxis and swimming velocity in Chlamydomonas

We previously showed that lack of tubulin glycylation impairs axonemal dynein regulation and ciliary motility in mouse sperm cells (22). Our immuno-cryo-ET experiments reveal a direct interaction of the microtubule binding domain of axonemal dyneins with glycylated tubulin sites (Figure 4 A). Thus, we checked if the lack of tubulin glycylation affects the swimming of Chlamydomonas cells. The ttll3-mutant from the CLiP library was an inappropriate background strain for swimming experiments, as the background strain itself shows significantly reduced swimming velocities compared to wild type (CC-124). Therefore, we generated a suitable CRISPR-Cas mutant ttll3-CRISPR from CC-124, which lacked tubulin glycylation in its axoneme (Supplementary Figure 7 A). Ttll3-CRISPR cells showed a slower phototactic response that formed an altered pattern (Figure 4 B and Supplementary Figure 8 A) and did not manage to accumulate cells at one side of the plate even after long times of exposure to light (Supplementary Figure 8 B). We then recorded high-speed movies of Chlamydomonas cells in liquid culture, which allowed us to accurately measure cell position, beating frequency and swimming speed. Surprisingly, ttll3-CRISPR cells swam faster than wild type (Figure 4 C, Supplementary Figure 7 B). Although the beat frequency was lower for ttll3-CRISPR cell when compared to wild type (Figure 4 D), ttll3-CRISPR cells showed higher forward displacement per beat cycle (Figure 4 E), explaining why ttll3-CRISPR swims faster than wild type. These results now show that the phenotype in Chlamydomonas and the previously observed phenotype of ttll3,8-/- mice (22) is mediated by the loss of a direct physical interaction between tubulin glycylation and the axonemal dynein microtubule binding domain.

**Figure 4:**
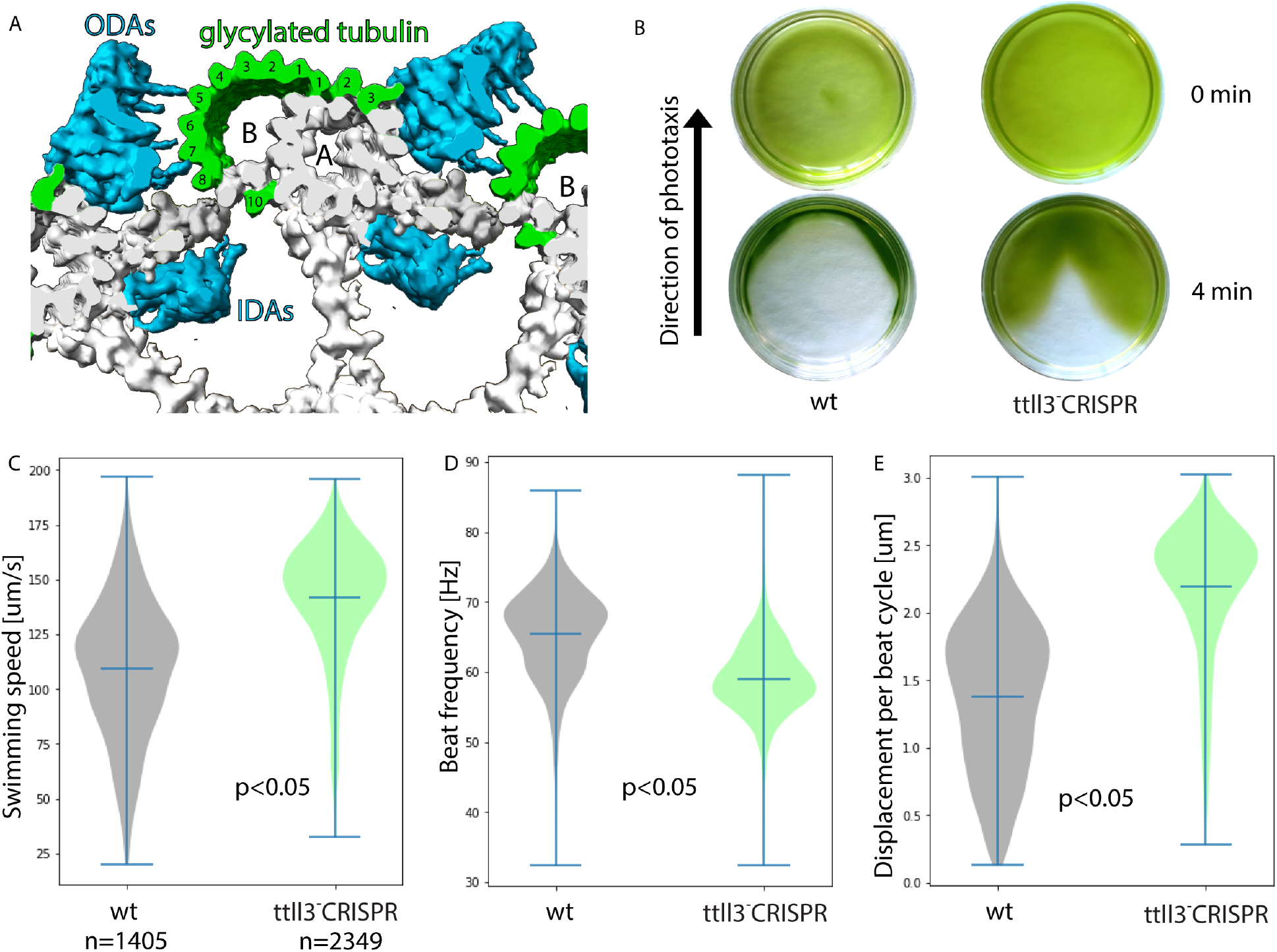
Tubulin glycylation is required for normal cell swimming behavior. **A**: 3D electron density model of the Chlamydomonas 96-nm repeat showing the interaction between ODAs and IDAs with glycylated tubulin. **B**: Phototactic response (negative) of wt and *ttll3^-^*CRISPR cells after 4 minutes of illumination with green light. Note that the pattern of phototaxis differs between wild type and *ttll3^-^*CRISPR. **C**: Violin plots of swimming speed distributions in wild type and *ttll3^-^*CRISPR cells. **D**: Violin plots of beating frequency distributions in wild type and *ttll3^-^*CRISPR cells. **E**: Violin plots of mean total displacement per beat cycle distributions in wild type and *ttll3^-^*CRISPR cells. P-values were calculated in python using a T-test (scipy.stats.ttest_ind).

### Axonemal integrity during the ciliary beat cycle is guaranteed by sliding of the NDRC over the polyglutamylated protofilament

Another axonemal component that directly interacts with the B-tubule is the nexin-dynein regulatory complex (NDRC), which is known to passively bridge neighboring microtubule doublets and contribute to the regulation of axonemal dyneins (29, 30). The NDRC specifically interacts with protofilament B9 (Figure 5 A), which we showed to be highly enriched in tubulin polyglutamylation. This is consistent with studies that link the interaction between the NDRC and polyglutamylation to axonemal integrity during ciliary beating (29,30). While our analysis showed a continuous decoration (every tubulin dimer) of the B9 protofilament by polyglutamylation, a previous study proposed that there is a single site for tubulin polyglutamylation per 96-nm repeat (27). This polyglutamylated site would provide a punctiform distribution of negative charges that would function as an electrostatic anchor for the interaction with the positively charged lobular region of the NDRC (31). In this scenario, and because doublets have been shown to slide during ciliary beating (32), it was proposed that the NDRC would stretch and/or tilt along the axoneme’s long axis (33) to preserve the interaction of its lobular region with the neighboring microtubule doublets and thus the stability of the axonemal structure. In order to test these hypotheses, we performed cryo-electron tomography on reactivated axonemes to simultaneously and precisely quantify inter-microtubule sliding (Figure 5 B) and visualize the speculated conformational change of the NDRC structure during microtubule sliding (Figure 5 C). Cumulative sliding displacements between neighboring doublets reached values of around 80nm measured over 1.5µm of ciliary length upon bending at physiologically relevant curvatures (Figure 5 B). Thus, we estimated that in distal sections of the axoneme, a single 96-nm repeat interacts with at least 3 contiguous axonemal repeats from neighboring doublets throughout a breaststroke beating cycle. We then analyzed the structure of NDRC complexes subjected to positive, negative and no sliding by sub-tomogram averaging. The NDRC did not deform nor tilt along the microtubule axis upon inter-microtubule doublet sliding (Figure 5 C). We conclude that, in order to preserve the interaction between the NDRC and the neighboring doublet during the ciliary beat cycle, the positively charged lobular region of the NDRC slides over the continuous distribution of negative charges generated by tubulin polyglutamylation all along protofilament B9 (Figure 5 A, right panel). These results change the way the interaction between the NDRC and microtubules is understood, providing a new framework for physical modeling of ciliary beating and its regulation.

**Figure 5:**
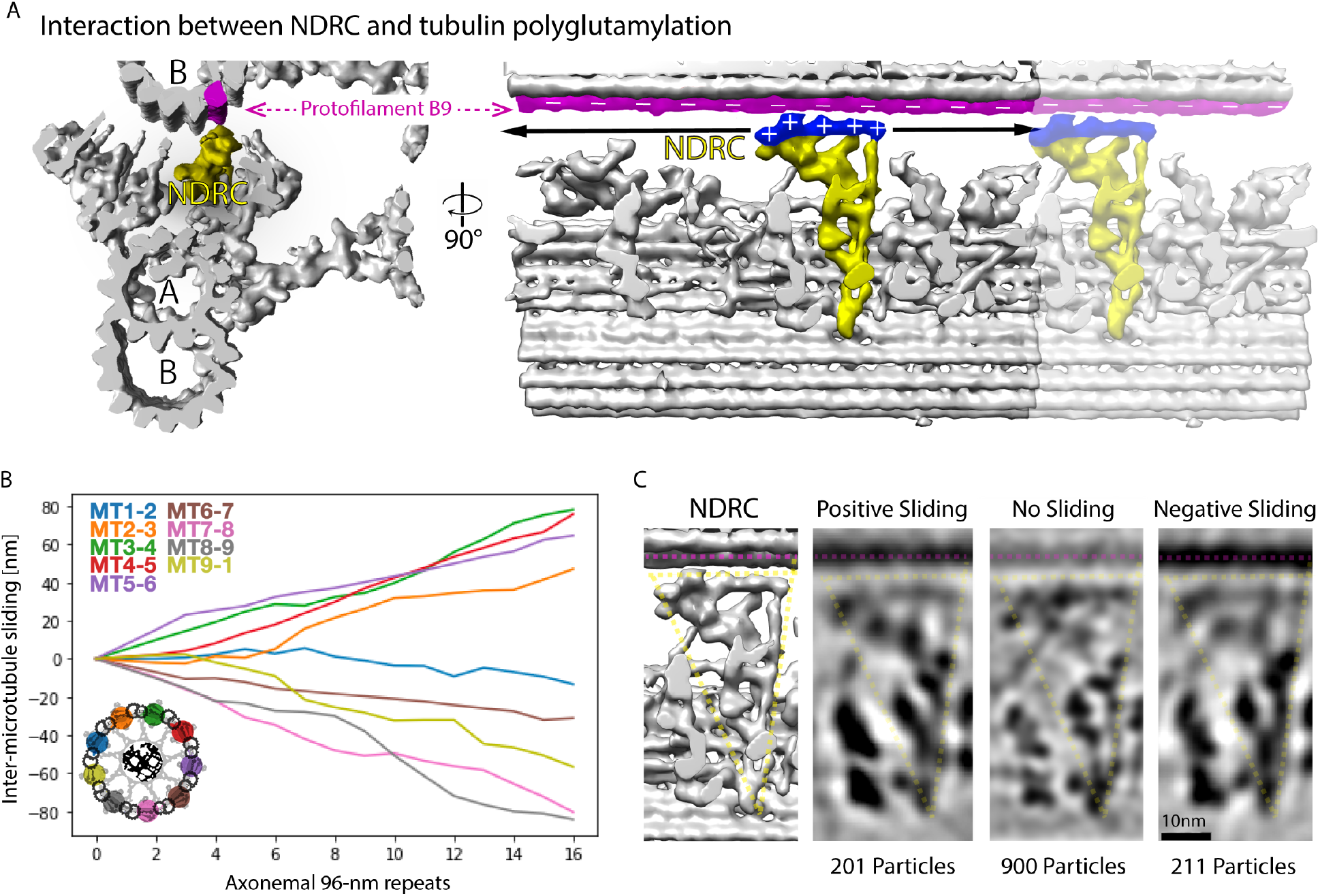
Intermicrotubule doublet sliding quantification and characterization of NDRC structure during ciliary bending. **A**: Electron density models showing the interaction between the NDRC (yellow) and protofilament B9 (magenta) in wild type cells. **B**: Measurement of the inter-microtubule sliding distributions of a bent axoneme. **C**: Slices through sub-tomogram average models of the NDRC subjected to different sliding conditions showing that the NDRC does not tilt nor stretch during ciliary bending *in vitro*.

## Discussion

With this work, we show that a combination of immunolabeling and cryo-electron tomography can reveal the nanometric location of tPTMs within higher order microtubule structures in their native context. Using this method, we show that the Tubulin Code forms a protofilament-specific nanopattern in cilia that reflects the molecular organization of axonemal structures and contributes to their regulation.

The mutually exclusive nanopatterning of axonemal tubulin polyglutamylation and glycylation is striking. The distribution of glycylation allows it to directly interact with the microtubule binding domains of dynein arms. Dynein arms are known to be the main driver of the beating activity of cilia, contributing to its amplitude, frequency, and waveform (reviewed in 6, 34). We and others have shown that the lack of tubulin glycylation can affect the motile function of cilia in zebrafish and mice (17, 22, 35). Consistent with this, our experiments in Chlamydomonas show an effect on ciliary based motility upon depletion of tubulin glycylation: mutant cells swam faster than wild type cells, although their beat frequency remained normal. The fact that these cells advance more with every beat cycle suggests that they have a stronger powerstroke, which might be caused by an altered interaction between the microtubule binding domain of dyneins and the microtubular surface. Although faster, ttll3-CRISPR cells appear to have an impaired fine control over their movement as their phototactic response is affected by the lack of glycylation. These results indicate that the described tubulin glycylation distribution is required for the regulation of the activity of axonemal dyneins through a direct physical interaction with their microtubule binding domains.

Our data in motile cilia show that depletion of tubulin polyglutamylation or glycylation does not disrupt axonemal assembly in Chlamydomonas, indicating that these modifications do not play a major role in the incorporation of axonemal components onto the algae ciliary microtubules. Nevertheless, tubulin polyglutamylation is important for axonemal integrity and proper ciliary motility through its interaction with the NDRC (29, 30, 36). The electrostatic interaction between positively charged residues in the DRC4 protein N-terminus (31) and negatively charged glutamates from polyglutamylated tubulin C-termini has been shown to contribute to the physical integrity of the axoneme (27, 37).

According to the polyglutamylation distribution previously proposed (27), the NDRC would have to act as a linear spring element, rigidly anchored at both ends (revised in 33), that tilts or stretches during ciliary beating to accommodate microtubule sliding. However, our results indicate that the NDRC does not act as a simple linear elastic element. Instead, the positively charged globular region of the NDRC glides over the negatively charged B9 protofilament from the neighboring microtubule. We propose that the interaction between the NDRC and polyglutamylated tubulin acts as a sliding joint which ensures a proper lateral interaction between microtubules throughout a whole ciliary beating cycle. This sliding joint energetically favors microtubule motions parallel to the ciliary axis while it prevents their relative displacement perpendicular to the ciliary axis.

The model described here proposes that tubulin polyglutamylation is required for proper ciliary beating because it couples the dynamics of microtubule sliding and microtubule bending through its interaction with the NDRC, which in turn participates in dynein regulation. Changes in intraciliary calcium concentration, which are known to influence beating frequency and wave form (38), could lead to a modulation of the electrostatic interaction of the NDRC with the neighboring microtubule doublet. These results suggest the existence of a flagellar beat control mechanism by which inter-microtubule sliding force generation can be locally modulated by the electromechanical interaction between the NDRC and the B-tubule, providing a new and alternative answer to the problem of ciliary beat regulation. The evolutionarily conserved nature of the NDRC and of the polyglutamylation pattern can explain the motility phenotypes found in Chlamydomonas (39) and mice (18, 22).

This study demonstrates that a specific polyglutamylation/glycylation pattern is correlated with the position and regulation of axonemal components, namely dynein arms and the NDRC. The question that naturally arises is how the modifying enzymes are able to generate such fine patterns. The enzymes involved in tubulin polyglutamylation and glycylation are assumed to compete for their initiation sites on glutamate residues present in the tubulin C-terminus (40, 41). Our results strongly suggest that the modifying enzymes for tubulin polyglutamylation and glycylation specifically recognize their targets and shed some light on the sequence of events giving rise to the precise nanopatterning of tPTMs. We found that the distal tip of both steady state and growing flagella is enriched in unmodified tubulin. The very distal part of the axoneme is known to undergo a continuous tubulin turnover also in steady state cilia (42). Thus, the absence of tPTMs in this area might allow for a faster exchange of tubulin dimers at the plus-end. Polyglutamylated tubulin signals remain at constant levels along the cilium during assembly, suggesting that most of the polyglutamylases rapidly carry out their activity in the tip area, apparently thread-milling with the growing B-tubule tip. On the other hand, tubulin glycylation accumulates over time along the cilium, indicating a less localized activity of diffusive modifying enzymes. Our data suggest that acetylation is applied to the assembling microtubules before glutamylation, and that glycylation is the last among these 3 modifications to be performed. Thus, we could infer that protofilament B9 is made unavailable for the TTLL3 enzyme (glycylation) by the early activity of TTLL9 enzyme (polyglutamylation). Experiments using fluorescently labeled versions of the enzymes involved will be useful to clarify their mode of action during ciliary assembly.

It is possible that TTLL9 recognizes its specific target by site-specific cues that are already present in the assembled microtubule doublets, presumably patterns of microtubule internal proteins or, alternatively, differential protofilament binding affinities defined by tubulin lattice curvature. The presence of FAP52 and/or FAP90 proteins along protofilament B9 (9) could explain the protofilament-specific activity of modifying enzymes during ciliary assembly. In particular, FAP90 occupies areas in the extraluminal side of the microtubule around protofilament B9, potentially serving as a cue for specific activity of polyglutamylation enzymes. It will be interesting to study whether the lack of these MIPs interferes with the modification of axonemal microtubules. Experiments on the step-by-step assembly and maturation of the axoneme during ciliogenesis might be required to ultimately clarify the sequence of events giving rise to this precise biochemical nanopatterning.

With this study we reveal that the Tubulin Code of motile cilia consists of a protofilament specific nanopattern of tPTMs that is consistent with the organization of the axoneme and the distribution of specific functional interactions between axonemal components and the microtubule. This provides a framework to mechanistically understand the specific role of different tPTMs for the regulation of ciliary components that are required for proper ciliary beating. Here we focused on the mutually exclusive nanopattern glycylation and polyglutamylation, but the same approach can be used to investigate other axonemal tPTMs and their potential interactions with other ciliary components, such as the intraflagellar transport machinery (IFT). Similarly, this new understanding of the Tubulin Code at the nanoscale will revolutionize the way we understand the function of tPTMs in other stable microtubular structures, such as centrioles and cortical microtubules.

## MATERIALS AND METHODS

### Chlamydomonas reinhardtii Liquid Culture

For this study we used the following cell strains: Wild type(+) (strain CC-125, N.H. Gillham 1968, Pröschold et al. 2005); ttll3-(-) strain (CLiP 124984 (26)); ttll3-CRISPR(-) strain (CRISPR-Cas mutant); tpg1(-) strain (CC-5245,Tomohiro Kubo, George Witman lab, University of Massachusetts Medical School, May 2016); ida5(+) strain (CC-3420, Ritsu Kamiya, University of Tokyo, July 1997). For each liquid culture, approximately 300μl of cells were transferred under sterile conditions from solid agar-TAP plates to 300ml of liquid TAP and dispersed by vigorous shaking. Cells were grown for four days with continuous aeration under a 14/10hr light/dark cycle at 23°C.

### Axonemal isolation from Chlamydomonas

300ml of 4 days old Chlamydomonas culture in liquid TAP were used for each isolation experiment. After centrifugation at 1200g (JLA16.25) for 10 minutes at room temperature (slow break), cells were resuspended in 10mL of chilled 10mM HEPES. Clean cell suspensions were centrifuged at 1500g (rotor JA25.50) for 10 minutes at 4°C (no brake). In order to deciliate the cells, they were resuspended in 10mL of 5mM Dibucaine in chilled HDMS and incubated for 2 minutes on ice. Immediately, 10mL of HDMS containing 1mM DTT, 1μM Aprotinin and 0.5mM EGTA were added. Cell bodies were removed by centrifugation at 1500g (rotor JA25.50) for 20 minutes at 4°C (no brake). The supernatant was centrifuged at 30000g (rotor JA25.50) for 15 minutes at 4°C (max. brake). The resulting flagellar pellet was resuspended in 2ml of HMDEK buffer with 1mM DTT, 1uM Aprotinin and 0.1mM PMSF. The remaining cell debris was removed by centrifugation at 800g for 3 minutes at 4°C (Ependorf, 5415R tabletop centrifuge). Clean flagellar suspensions were demembranated by addition of 1% Igepal and incubation on ice for 20 minutes.

### Axonemal isolation from mouse trachea

Fresh 1ml stock 2mg/ml Digitonin solution was previously prepared by heating up to 95°C until the detergent dissolves. Dissected tracheas were obtained in cold PBS at 4°C. Tracheas were immersed into 2ml of deciliation buffer (10mM Tris, 50mM NaCl, 10mM CaCl2, 1mM EDTA, pH adjusted to 7.5 using HCl at 4°C, containing 20μM Digitonin, 7mM β-mercaptoethanol, 1% Protease inhibitor cocktail Sigma Aldrich, P1860). The buffer was gently flushed 20 times through each trachea to boost deciliation for about 5 minutes. Clear supernatant was then collected and centrifuged at 500g for 1 minute at 4°C. The resulting supernatant was collected and centrifuged at 5000g for 5 minutes at 4°C to obtain a ciliary pellet. The pellet was resuspended in 20μl of HMDEK buffer containing 1mM DTT, 1uM Aprotinin, 0.1mM PMSF and 1%Igepal.

### Western blot

Wild-type (CC125), ttll3- (CLiP 124984) and ttll3-CRISPR strains were cultured for 4 days in liquid TAP as described above. Flagellar suspensions obtained through isolation following the described protocol were loaded in ready-to-use 4-12% electrophoresis gels (NuPAGE Bis-Tris Mini, Invitrogen), together with a molecular weight ladder (PAGERuler Plus or Chameleon Duo). Electrophoresis was carried out in MOPS buffer by applying a constant current of 50mA for 1h (Electrophoresis kit, Invitrogen). Tubulin concentrations were assessed either by using Coomassie staining or acetylated tubulin content. A second gel was loaded with matching sample concentrations. The resulting bands were transferred onto a membrane using the iBlot machine (iBlot 2 NC Mini Stacks) with a 3 steps program of 20V each (10min approx total run time). After washing it with deionised water, the blotted membrane was automatically immunolabeled using an iBind machine (using the iBind Flex FD solution kit) with a primary rabbit antibody (1:10000 dilution) against glycylated tubulin (Glypep1) and (in the case of ttll3-CRISPR) a primary mouse antibody against acetylated tubulin. Secondary antibodies consisted of infrared-fluorophore-conjugated antibody (1:3000 dilution) against rabbit (800 CW) and, in the case of ttll3-CRISPR, an additional secondary (1:4000) antibody against mouse (680LT). Using a LI-COR fluorescence scanner and Image Studio software, fluorescence images of the tubulin glycylation and acetylation associated channels were acquired and analyzed.

### Immunolabeling of axonemes for cryo-ET

Chlamydomonas axonemal suspensions were diluted to a very low concentration to expose isolated axonemes to saturating antibody concentrations. Optimal dilution was achieved when, after quickly loading 4μL of dilution on a glass slide and covering it with a coverslip, approximately 15 axonemes appeared adhered to the glass surface per field of view (∼6650μm2, observation with a 60X oil immersion objective using phase contrast). 1 μL of antibodies against a particular tubulin post-translational modification were added to 100μL of diluted axonemal suspension and incubated for at least 1.5h inside 0.5mL tubes lying over an ice surface prior to plunge-freezing. In the case of mouse trachea axonemes, antibodies were used 1:100 directly on isolated axonemes. Antibodies used:

polyE: glutamate chains (>4E), AG-25B-0030 Adpipogen
Glypep1: glycylated β-tubulin, Gadadhar et al. 2017

### Plunge freezing of Chlamydomonas and mouse cilia

A Leica Automatic Plunge Freezer EM GP machine was used to perform plunge freezing as it counts with an integrated humidity control chamber. Cu supported 3.5/1 Holey Carbon grids (Quantifoil) were glow discharged on both sides for six seconds each using atmospheric air in order to render them more hydrophilic. In the meantime, the plunge freezer chamber temperature was set to 32°C in order to obtain and maintain high humidity (above 80%). The plunge freezer was set to detect the wetting of the filter paper (Whatman) upon blotting (around 2 seconds blotting time, no automatic plunging), which was mounted on one side only with the purpose of blotting from the grids back side. The sample was loaded on both sides (3.5μL total) and incubated for 30 seconds before the addition of 1.5μL of sonicated gold fiducials suspended in PBS, followed by immediate blotting. In many cases automatic blotting was not optimal, requiring swift manual blotting before plunging. Grids were stored in custom made grid-boxes (MPI-CBG Workshop) which were themselves stored in cryogenic tanks.

### Cryo-Electron Tomography

Acquisition of cryo-EM images were done with a 300 kV Thermo Fisher Titan Halo TEM with a field emission gun (FEG) electron source and a Gatan K2 Summit direct electron detector with energy filter using a slit width of 20 eV. To facilitate the location of regions of interest containing well preserved cilia, SerialEM (43) was used to generate a full grid overview by automatically acquiring and stitching low magnification (210x) images. Tilt series were acquired with SerialEM on areas of interest at 30,000x nominal image magnification, resulting in a calibrated pixel size of 4.72 Å (counted mode). Tilt series were recorded with 2° increments with a bidirectional tilt scheme from –24° to 42° and from -26° to –42°. The tilt step was varied up to 6° whenever dose limitations applied. The defocus target varied from –1.5 to -5.5μm and the cumulative dose was 95-120 e - per Å 2 per tomogram. Images were acquired in the dose fractionation mode with 0.25 s frame time to a total of around 8 frames per tilt image. Drift was kept below 1nm/s. The frames were aligned using K2Align software, which is based on the MotionCorr algorithm (44). Tomogram reconstruction was performed using Etomo from IMOD v.4.10.11-α using weighted backprojection (45). Contrast transfer function curves were estimated with CTFPLOTTER and corrected by phase-flipping with the software CTFPHASEFLIP, both implemented in IMOD (46). Dose weighted filtering was also performed using the mtffilter command from the IMOD package. To enhance the contrast of macromolecular structures and facilitate visualization/interpretation of tomograms and sub-tomogram picking, individual images were treated as to synthetically remove gold fiducials and linearly binned by two prior to tomographic reconstruction. In some cases, a nonlinear anisotropic diffusion filter by IMOD (45) was applied on low-contrast tomograms acquired closer to focus.

### Sub-tomogram averaging

IMOD v.4.10.11-α and PEET v1.11.0 were used for sub-tomogram averaging (47). Using 3dmod, included in the IMOD package, tomograms were inspected looking for signs of adequate structural preservation, namely, roundness and straightness of the axonemes and presence of unaltered axonemal components such as radial spokes and dynein arms. To model the location of each 96-nm repeat within tomograms, the coordinates of points of the central axis of A-tubules at the base of every RS2 were determined manually and saved in .mod format (supported by IMOD software). Approximately, at least a thousand of these points were determined per experimental condition. Using PEET (v 1.11.0), each sub-tomogram containing a 96-nm repeat was aligned to a common reference so that all microtubule axes remained parallel to each other with the right polarity. The particles from each microtubule doublet were averaged independently to determine the angles of rotation that should be applied to each microtubule doublet to pre-align it with the common reference, providing a good starting point for automated fine alignment. Missing wedge compensation was used, one single particle was commonly used as an initial reference per experiment. Soft masks outlining the 96-nm repeat structure were used for alignment optimization. The maximum search ranges for translations and rotations of each particle during alignment to iteratively refined references were estimated based on the precision achieved during manual modeling and pre-alignment, normally resulting in values of +-12° around the Y axis, +-6° for the remaining axes and 8 pixels for translations in each direction. This strategy, laborious compared to fully automated sub-tomogram averaging routines, provides human-proofread averages without missing-wedge, chirality or polarity artifacts. Routinely, the final average was generated discarding the worst 10% of particles based on their cross-correlation coefficient score against the final reference (consisting of the average of the 66% best aligned particles). Visualization of averaged electron density maps was performed in 3dmod from IMOD. 3D rendering of isosurfaces and structure fitting was performed using UCSF Chimera (v 1.14) (48). Thresholds for signal detection in electron density maps were calculated using False Discovery Rate Control software (49).

### Ultrastructural expansion microscopy of regrowing flagella

Chlamydomonas cells from the strain CC125 were inoculated from a TAP agar plate to a liquid culture containing 400ml TAP, in a 500ml bottle, equipped with a bubbling device connected to compressed air. After growing in a light-dark cycle for 3 days, 200ml of culture was spun down at 500g for 2 minutes and the cell pellet resuspended in 40ml of 20mM HEPES, pH 7.2. The pH was lowered to 4.5 with acetic acid and quickly raised to 7.2 with NaOH. Cells were pelleted by centrifugation at 500g for 2 minutes and resuspended in TAP. The time zero of the time course during flagellar regrowth is when cells are fully resuspended. Cells were placed in a 50ml glass beaker and mixed with a magnetic stir bar at low speed. Cells were harvested every 5 minutes until time point 30 minutes, after which cells were harvested every 10 minutes until time point 60 minutes. One additional time point was collected at 90 minutes. At each time point 200ul cells were applied to a 12mm coverslip previously treated with poly-D-lysine. After 5 minutes the coverslip was lightly blotted on the side with a paper towel and placed in absolute methanol at -20°C. After 15 minutes the coverslip was equilibrated in 1x PBS for 15 minutes.

The coverslips were then treated for expansion microscopy (Ultrastructure expansion microscopy (U-ExM); Davide Gambarotto, Virginie Hamel, Paul Guichard; Methods in Cell Biology, ISSN 0091-679X). The coverslips were submerged into cross linking prevention solution (2% acrylamide (Sigma, A4058) and 1.4% formaldehyde (Sigma, F8775)) for 3 hours at 37°C. Gel solution was prepared for each coverslip as follows: Ice cold monomer solution (sodium acrylate (Sigma, 408220) 21.123%,acrylamide 11.10%, N,N-methylenebisacrylamide (Sigma, M1533) 0.1% in 1x PBS) was mixed with TEMED and APS, vortexed, and 35ul of the final gel solution were placed on a slice of parafilm, ice cold, and the coverslip layered on top with the cells facing the gel drop. This was repeated for all the coverslips of all the time points. Coverslips were placed in a sealed humid chamber and incubated 15 minutes on ice and 1 hour at 37°C. Cover slips were then submerged in a denaturing solution (200mM SDS, 200mM NaCl, 50mM Tris-BASE) for 15 minutes at room temperature. Gels were detached from the coverslips and placed in microfuge tubes and heated at 95°C for 1 hour and 30 minutes. After denaturing, the excess of denaturing solution was removed, and gels were expanded by incubating them 2 times in 150ml of distilled water for 30 minutes each and 1 time overnight. The following day a quarter of a gel of the time point of interest was stained as follows. The antibodies used will be listed below. The gels of interest were shrunk by submerging them in 1x PBS for 20 minutes, 3 times. Primary antibodies were mixed at 1:300 dilution in 2% BSA solubilized in 1x PBS and incubated for 3 hours at 37°C on a rocking platform. Primary antibody incubation was followed by 3 washes with 1x PBS added with Tween20 at 0.1% final concentration. Secondary antibodies were incubated for 3 hours at 37°C and then washed 3 times with 1x PBS added with Tween20 at 0.1% final concentration. Gels were expanded by placing them in distilled water for 20 minutes, for 2 times, and overnight for the last step. Stained gels were imaged with a Zeiss LSM980 in Airyscan mode, with a 63x Zeiss oil immersion lens with 1.46NA, with a 0.4% laser power and with Z stacks with 0.150um sampling rate.

Primary antibody list:

Anti polyG in rabbit; Adipogen, AG-25B-0034-C100
Anti PolyE in Rabbit; Adipogen, AG-25B-0030-C050
Anti alpha and beta Tubulin in goat; Absolute Antibody, F2C S11B
Anti acetylated tubulin in mouse; Sigma, T7451

Anti Centrin in mouse; kind gift from the Rosenbaum laboratory (Molecular cloning and expression of flagellar radial spoke and dynein genes of Chlamydomonas. B D Williams, D R Mitchell, J L Rosenbaum. J Cell Biol. 1986 Jul;103(1):1-11. doi: 10.1083/jcb.103.1.1).

All secondary antibodies were from Thermofisher Scientific:

-Donkey anti-Mouse IgG (H+L) Highly Cross-Adsorbed Secondary Antibody, Alexa Fluor Plus 488, A32766
-Donkey anti-mouse IgG Alexa 546, A10036
-Donkey anti-Rabbit IgG Alexa 546, A10040
-Donkey anti-goat IgG Alexa 647, A21447

The quantification of normalized signal densities was carried out with Fiji. Z stacks were projected including all Z intervals using the maximum intensity project in Zen Blue from Zeiss. The signal from tubulin was used to draw ROIs around the cilium of interest and the respective basal body. The intensity (measured as RawIntDen in Fiji) of the ROI was divided by its surface. The resulting intensity/surface of the cilium was divided by the intensity/surface of the respective basal body. This measurement is referred to as normalized intensity. Since polyglutamylation stains the basal bodies, the same primary, and therefore secondary, has been used to measure the intensities of the anti poly-glutamylation signal for both cilia and basal bodies. To normalize the signal for the glycylation, the gels have been labeled with antibodies against centrin that show specificity for the basal body area only, and not for the cilium. Since the antibody against centrin was raised in mouse and the antibody against glycylation was raised in rabbit, secondaries labeled with Alexa 547 and against mouse and rabbit were used.

### Generation of ttll3-mutants by CRISPR

The ttll3-CRISPR line was generated using CRISPR targeted insertional mutagenesis of CC-124 32M which was received as a kind gift from Susan Dutcher. A suitable CRISPR site (GTGCACTGTTGACCACGACGCGG) was chosen on exon two of ttll3-. An insertional cassette was generated by flanking a Blasticidin S Deaminase gene under the control of a RbcS2 promoter terminator pair with 50bp long homology arms upstream and downstream of the chosen cut site respectively by Gibson cloning. The insertional cassette was prepared for transformation by digestion with BspQI, followed by column purification. RNPs were formed by annealing guide crRNA with tracrRNA at 98C for 2 minutes at 50µm of which 2µL was added with 2µL Cas9 (at 10 µg/µL) to 16µL duplex buffer. Chlamydomonas cells, grown for 4-5 days on plates, were washed with gamete autolysin for three rounds over a total of 4 hours and heat shocked at 40°C during the last round. Cells were subsequently washed three times in TAP + 40mM sucrose (TAPS). ∼5×106 cells in 50µL were mixed with 1.5µg of the insertional cassette and 5uL of RNP solution, then electroporated in 10µL reactions using a Neon electroporator for 3 pulses of 12ms at 2300V. Electroporated reactions were dispensed into 1mL TAPS, recovered overnight, and plated on 1.5% TAP agar plates containing 50µg/mL Blasticidin S. 48 of the resulting colonies were picked and screened by PCR with primers amplifying the entire insertion. Select clones were sequenced to verify a correct insertion.

### Chlamydomonas swimming experiments

Flow chambers were made by mounting a thin coverslip (0.17mm) over a glass slide using double sided sticky tape (Tesa 05338). Liquid culture cells grown as explained above were pipetted inside the flow chamber right before imaging. An inverted Nikon Ti2 microscope was used in diascopic imaging mode using a white light transilluminator lamp (Nikon Intensilight C-HGFI). Movies were acquired at 387.4 fps and 0.564 um/pixel using a 20x objective (Plan Apo lambda, NA=0.75, WD=1000um) and a PRIM 95B 25MM camera (Teledyne Photometrics). All movies were acquired focusing on the middle of the flow chamber. The acquired movies were analyzed using the TrackMate plugin in Fiji (50) which provided accurate sub-pixel localizations for each cell at each point in time. A custom Python script was used to compute swimming speeds, beat frequencies and forward and backward displacements per beat cycle.

### Chlamydomonas phototaxis experiments

Cells in light phase were resuspended in a phototaxis assay buffer (5 mM Hepes pH 7.4, 0.2 mM EGTA, 1 mM KCl, and 0.3 mM CaCl2, ∼107 cells/mL) and placed under red light for more than an hour (51). 3ml of liquid culture were placed on a plastic petri dish. The petri dish was illuminated from the side with a green LED light (60mW), inducing a negative phototactic response in the cells. The set-up was covered with a black box and imaged at different timepoints. A picture from the petri dish was taken with a smartphone.

### Reactivation of Chlamydomonas axonemes for MTd sliding measurement and NDRC structure characterization

Following axonemal isolation, 150μM ATP was added to the axonemal suspension. After inspection under the light microscope of a correct beating behavior of the axonemes in the mentioned condition, they were prepared for plunge freezing as described below. ATP was added just prior to plunging.

### Microtubule sliding quantification

To measure the deformations of the ciliary structure during active and passive beating, isolated Chlamydomonas axonemes were resuspended in HMDEK buffer containing 150μM ATP, and 1mM Casein. Both axonemal preparations were plunge-frozen as described in the Plunge freezing section for immunolabeled axonemes. Cryo-ET tilt series of axonemal sections presenting different curvatures were acquired at 24.000X magnification from -60 to 60 degrees. Tomographic reconstructions were carried out (see Cryo-ET), enabling manual annotation of each 96-nm repeat position in the axoneme and the discretization of the ciliary axis. A custom Python script was used to compute the orthogonal projection of each 96-nm repeat over the discretized ciliary axis, by finding the smallest distance between the localization of each repeat and the discretized ciliary axis. Then cumulative sliding displacements between each pair of neighboring microtubule doublets were calculated based on these projections. The values shown in Figure 3 have been filtered with a 5-element moving average to reduce noise and present overall displacement trends for each pair of microtubule doublets. The measured sliding distributions between different doublets were in agreement with the bending plane observed in tomograms. 96-nm repeat sub-tomograms associated with regions of negative (n=211), positive (n=201) and no sliding (n=900) were independently analyzed by sub-tomogram averaging (see Sub-tomogram averaging).

## Acknowledgments

We thank the Electron Microscopy and Light Microscopy facilities from both MPI-CBG Dresden and Human Technopole Milan for their instrumental support. We thank the Biomedical Services from MPI-CBG Dresden for providing the mice tissue used in this study. We thank Prof. Carsten Janke, Dr. Dennis Diener and Prof. Joel Rosenbaum and Prof. Susan Dutcher for sharing reagents. We thank Dr. Florian Jug, Dr. Iain Patten, Dr. Samuel Lacey, Dr. Helen Foster and Dr. Dennis Diener for comments on the manuscript. We thank Prof. Carsten Janke, Dr. Dennis Diener and Prof. Joel Rosenbaum for sharing antibodies.

## Author contributions

Conceptualization: GP, GAV

Methodology: GAV, NK, FM, APN, GP

Investigation: GAV, NK, FM, APN, GP

Visualization: GAV, NK, FM, APN, GP

Funding acquisition: GP, NK, APN

Project administration: GP

Supervision: GP

Writing – original draft: GAV, GP

Writing – review & editing: GAV, NK, FM, APN, GP

## Supplementary Figures

**Supplementary Figure 1:**
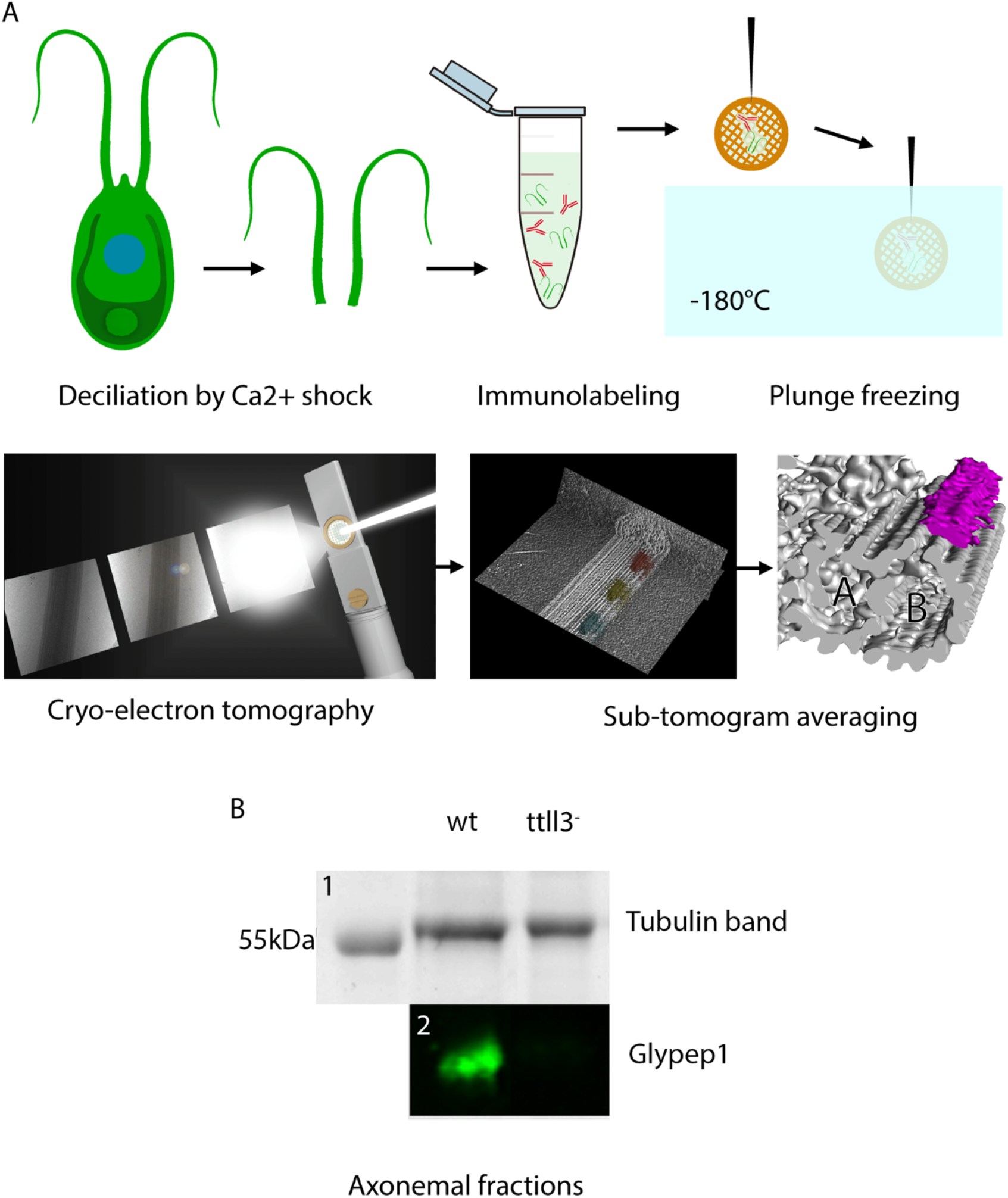
Immunolabeling method for cryo-electron tomography and sub-tomogram averaging. **A**: Chlamydomonas cells were deciliated using a calcium shock treatment and their axonemes were isolated as described in Materials and Methods. Axonemes were immunolabeled in suspension prior to their application to the EM grid. They were then plunge-frozen and imaged in cryo-electron tomography. Sub-tomogram averaging was performed resolving the nanometric location of bound antibodies in the context of the axonemal 96-nm repeat. **B**: Western Blot as a control for the phenotype of the CLiP Library mutant *ttll3^-^*. **1**: Total protein stain showing similar loadings for wt and *ttll3^-^* axonemal samples based on the signal from the tubulin band. **2**: Fluorescence Western Blot showing the lack of tubulin glycylation in *ttll3^-^* axonemes.

**Supplementary Figure 2:**
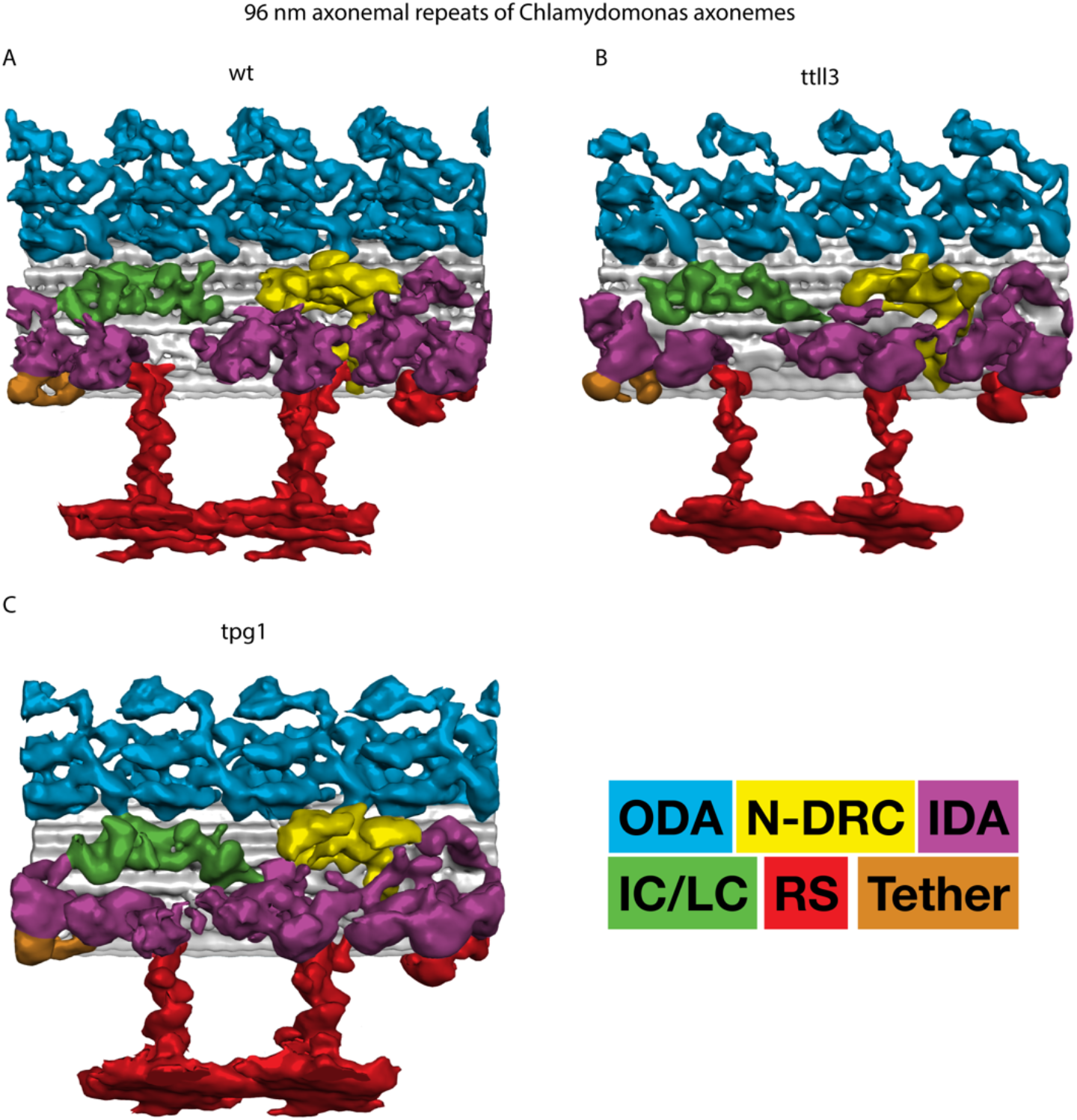
Knock-out of tubulin post-translational modifying enzymes does not alter the macromolecular structure of the Chlamydomonas axoneme. Electron density models of the axonemal 96-nm repeat from different Chlamydomonas strains generated by sub-tomogram averaging. **A**: wild type. **B**: ttll3-strain. **C**: tpg1 strain. Note that, at macromolecular level, the overall structure of the axonemal 96-nm repeat is complete.

**Supplementary Figure 3:**
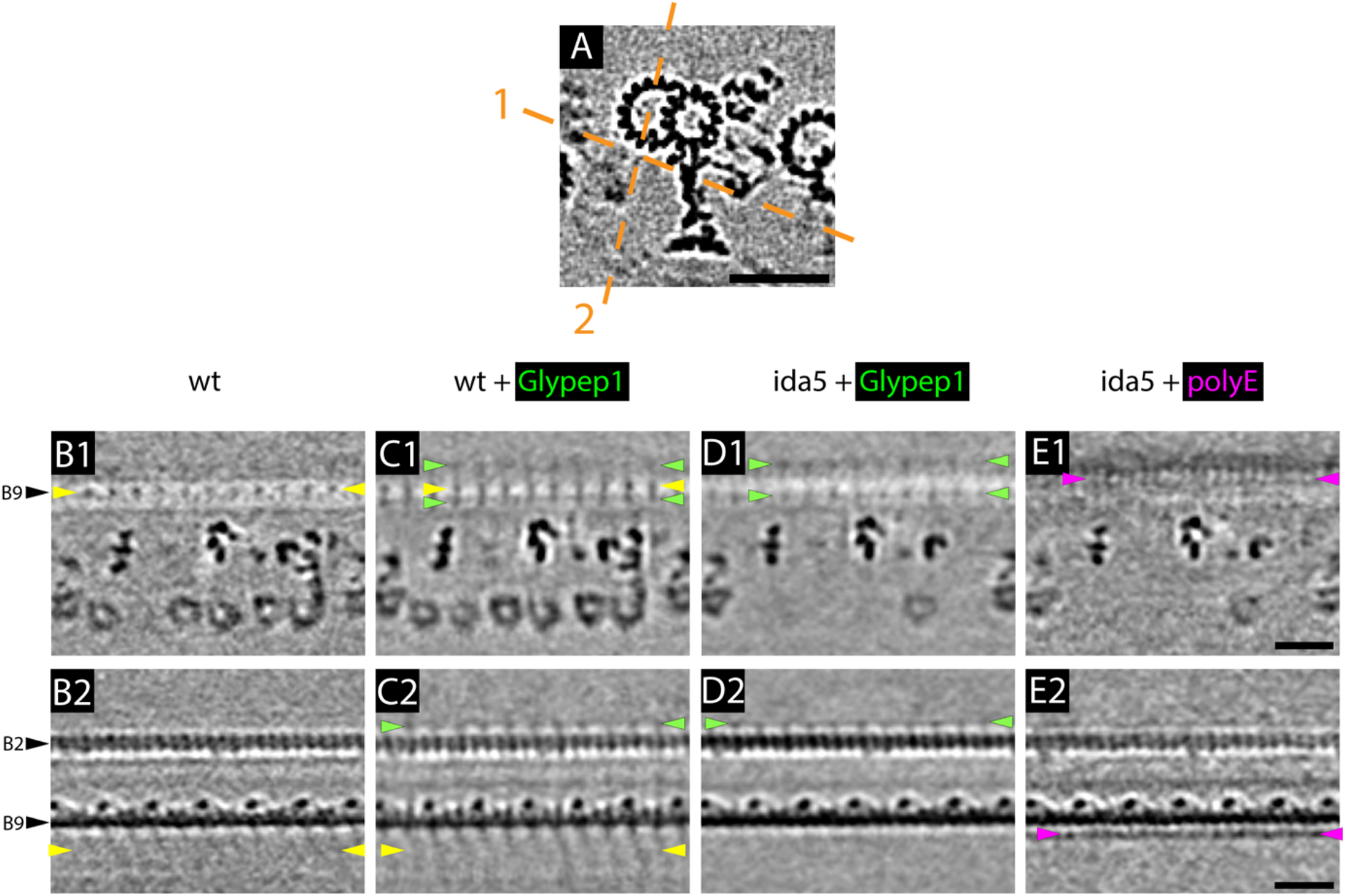
Tubulin glycylation and polyglutamylation form a complementary pattern in the Chlamydomonas axoneme. Slices through 3D electron density models of the Chlamydomonas axonemal 96-nm repeat generated by sub-tomogram averaging. Black arrowhead labels indicate the numbering of the different protofilaments. **A**: Cross-section of the axonemal 96-nm repeat. Orange dashed lines depict the orientation of the planes at which the different longitudinal sections were taken. **B**: Wild type. Note the presence of IDA stalks and microtubule binding domains indicated by yellow arrowheads. **C**: Wild type axonemes decorated with Glypep1 antibodies. Note the presence of extra densities indicating the binding site of antibodies (green arrowheads). **D**: Ida5 axonemes decorated with Glypep1 antibodies. Here, most IDAs are missing and it is possible to observe a lack of labeling over protofilament B9. **E**: Ida5 axonemes decorated with polyE antibodies. Note the presence of polyE immunolabeling densities specifically over protofilament B9 indicated by magenta arrowheads. Scale bars: 20nm.

**Supplementary Figure 4:**
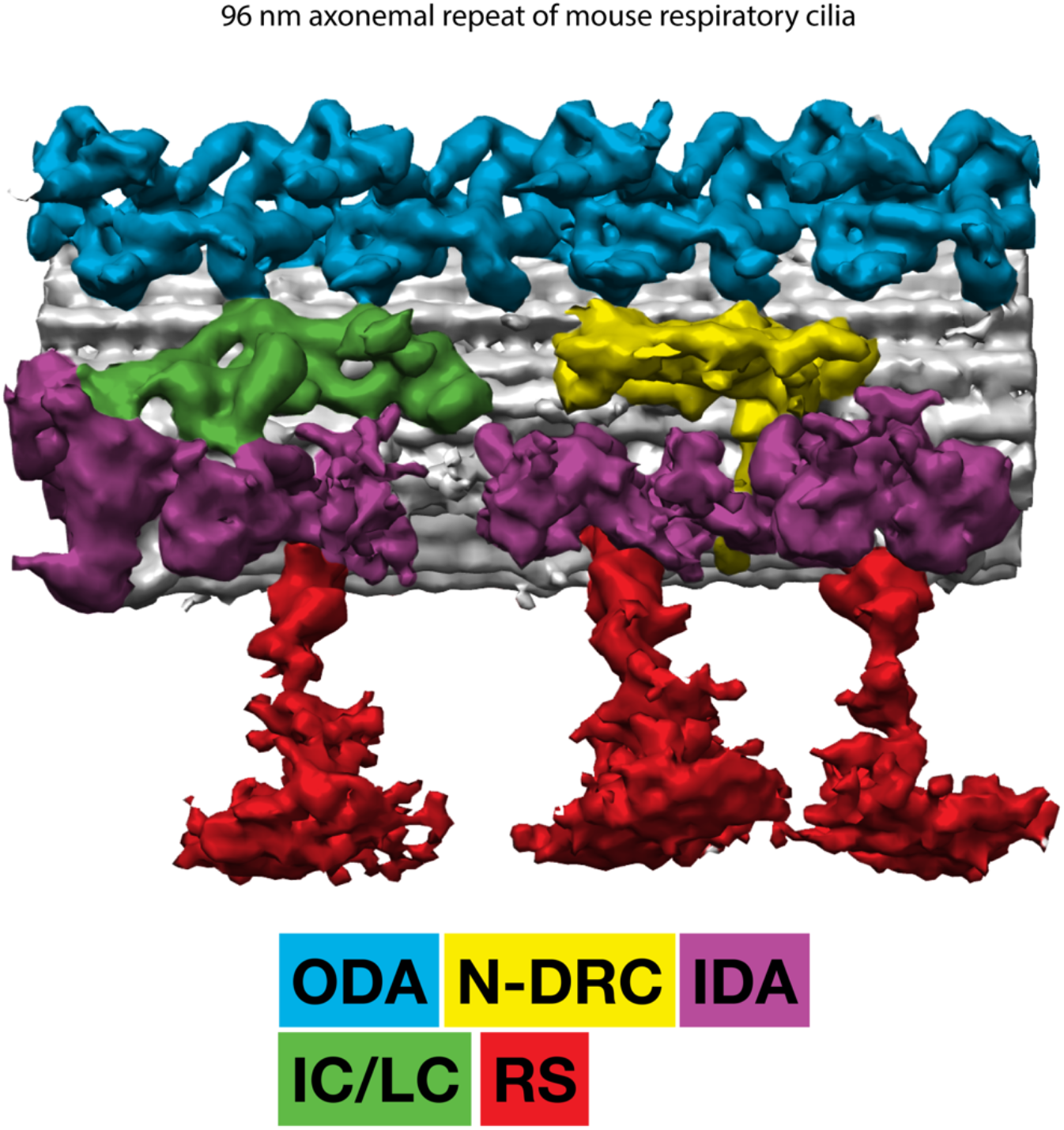
Isolation of mouse respiratory cilia successfully preserves the axonemal 96-nm repeat structure. Electron density model of the axoneme of mouse respiratory cilia generated by sub-tomogram averaging. All macromolecular components can be observed in the structure albeit a distortion of the radial spoke heads is present due to sub-optimal sample preservation.

**Supplementary Figure 5:**
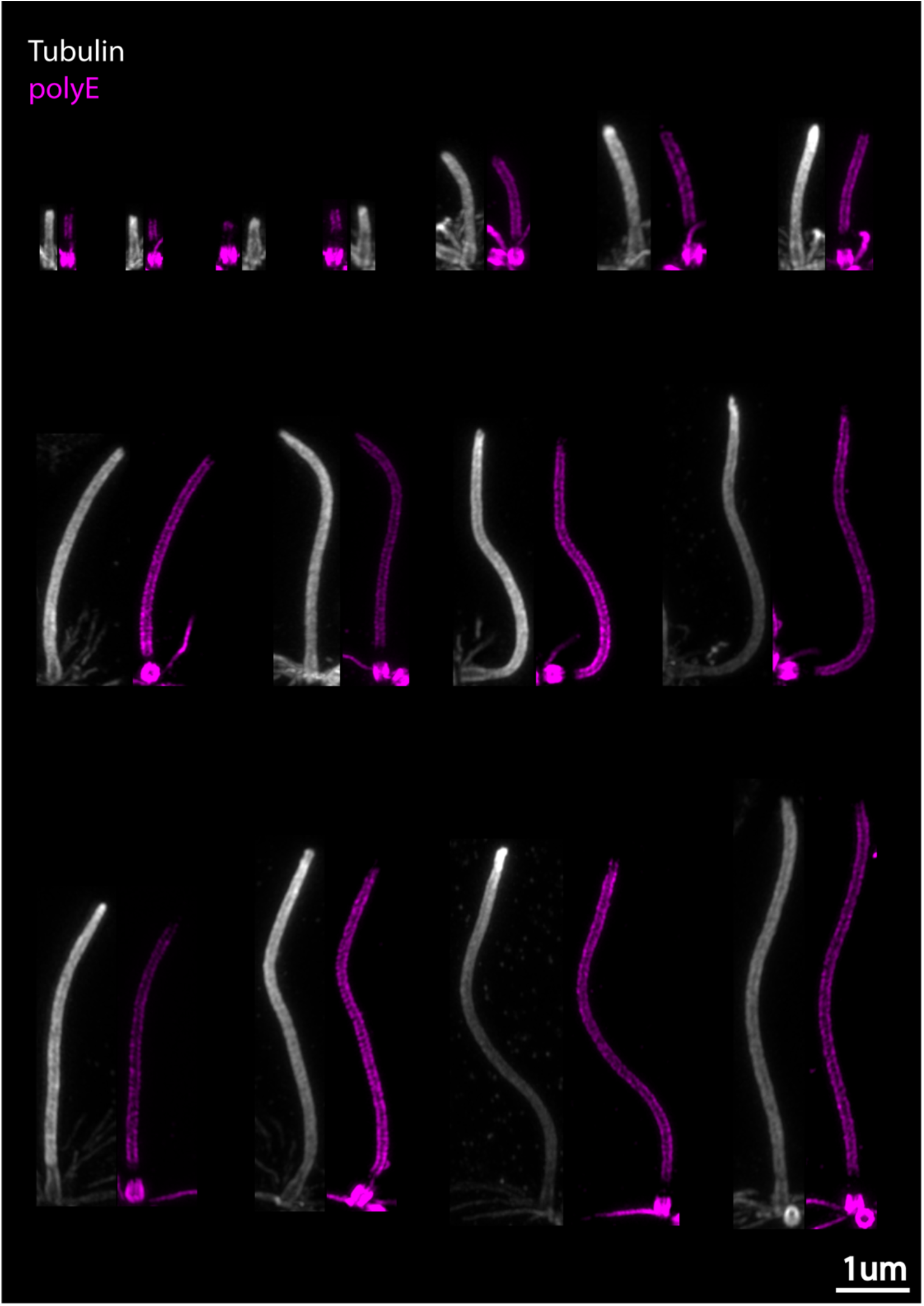
Tubulin polyglutamylation levels are constant along the axoneme during ciliary regrowth. Representative images from U-ExM experiments used to quantify the density of signal associated to tubulin polyglutamylation over the axoneme normalized to tubulin polyglutamylation levels in the basal body. Tubulin polyglutamylation is shown in magenta while tubulin is shown in gray. Scale bar: 1um.

**Supplementary Figure 6:**
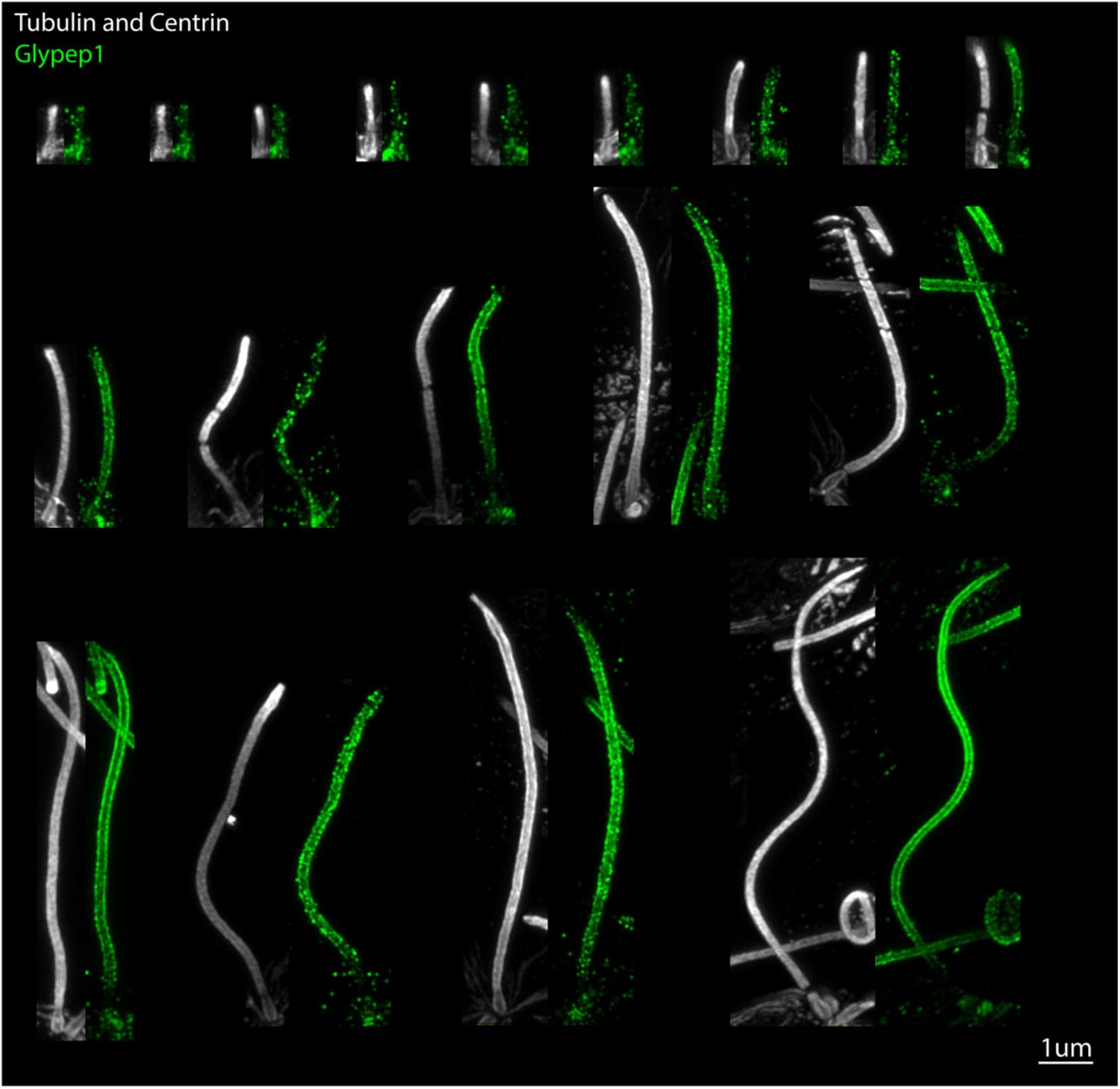
Tubulin glycylation slowly builds up in the axoneme during ciliary regrowth. Representative images from U-ExM experiments used to quantify the density of signal associated to tubulin glycylation over the axoneme normalized to centrin levels in the basal body. Tubulin and centrin labelings, sharing the same secondary antibody, are shown in gray while tubulin glycylation is shown in green. Note that for short axonemes the Glypep1 signal is sparsely localized along the axoneme while it gains density in longer flagella. Scale bar: 1um.

**Supplementary Figure 7:**
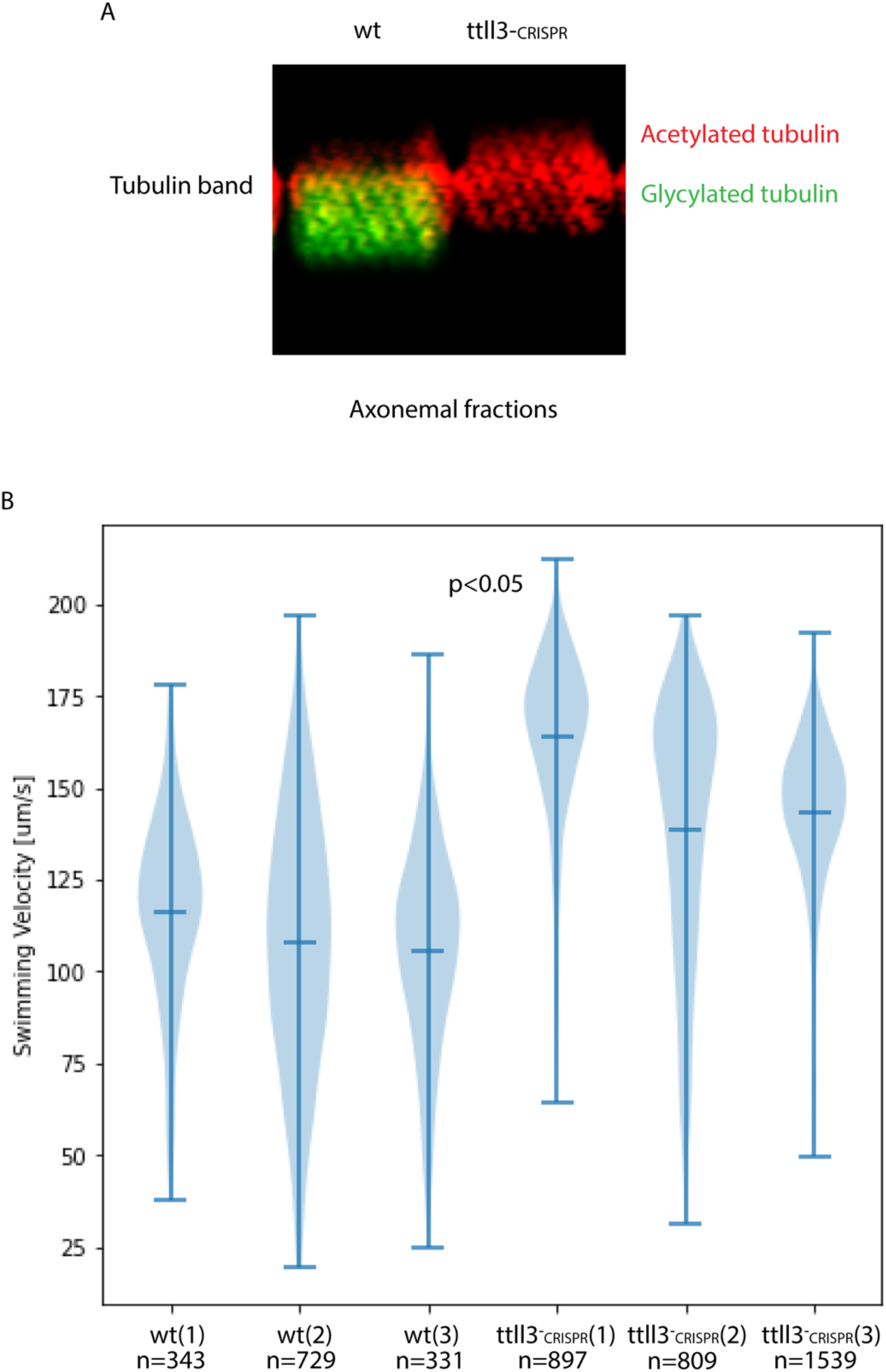
Lack of glycylation induces faster swimming in Chlamydomonas. **A**: Immunofluorescence image of a western blot membrane showing the lack of tubulin glycylation in the CRISPR-Cas mutant ttll3-CRISPR. Red: acetylated tubulin, Green: glycylated tubulin. **B**: Violin plot of three independent experiments measuring the swimming speed of both wild type and ttll3-CRISPR Chlamydomonas cells. P-values were calculated in python using a T-test (scipy.stats.ttest_ind).

**Supplementary Figure 8:**
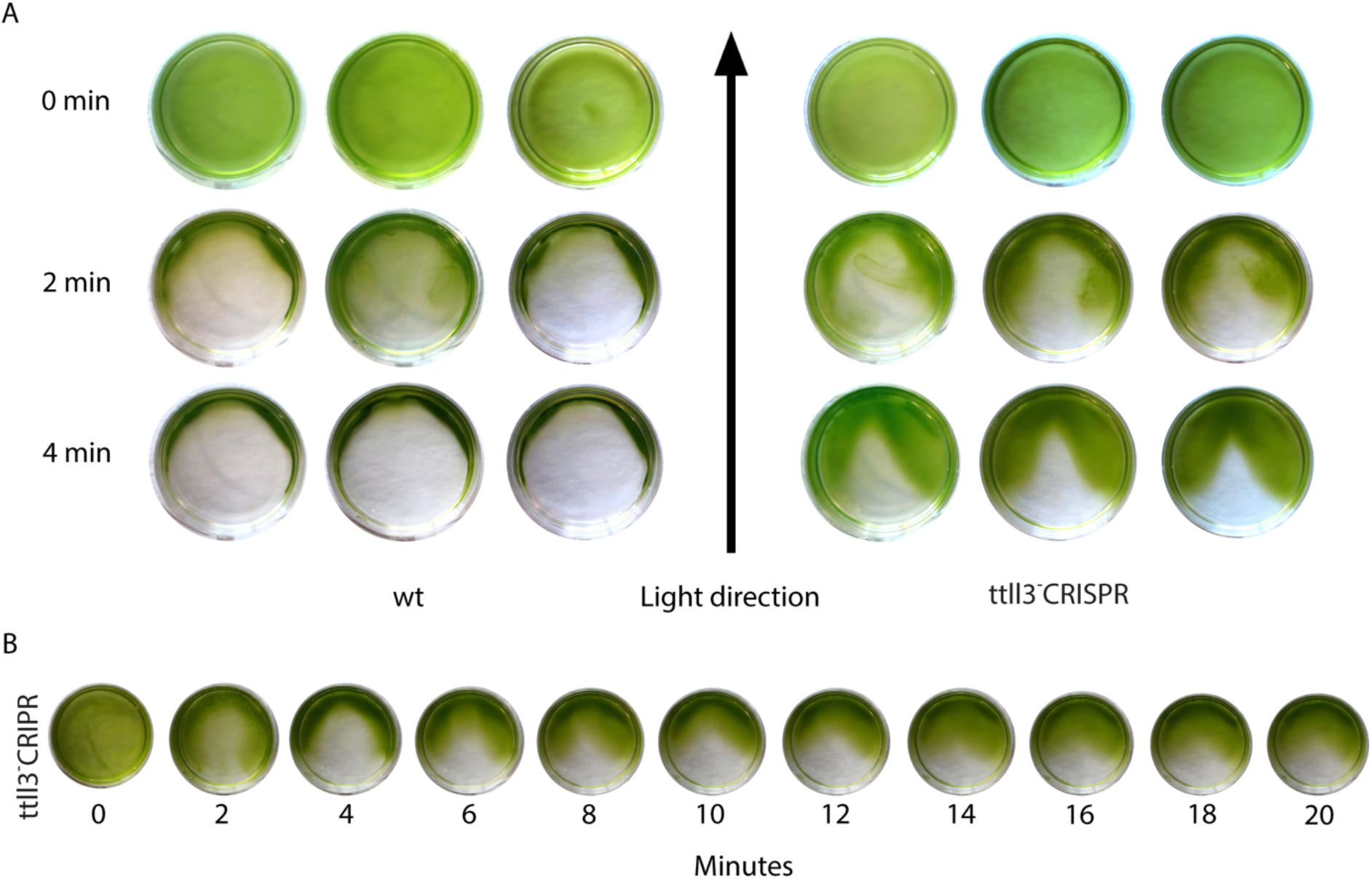
Lack of glycylation impairs phototaxis response in Chlamydomonas. **A**: Three replicates of a phototaxis experiment showing a normal phototaxis response of wild type cells and an impaired phototaxis in *ttll3^-^*CRISPR cells. **B**: Long term phototaxis assay with *ttll3^-^*CRISPR cells showing the lack of convergence of the cells after long exposures to light.

